## Supplementary Materials for "Light modulates task-dependent thalamo-cortical connectivity during an auditory attentional task"

**Supplementary Information Materials and Methods**

**Table S1: Peak coordinates of the left ROIs across subjects.**

|  | Thalamus |  |  | IPS |  |
| --- | --- | --- | --- | --- | --- |
| x | y | z | x | y | z |
| -14 | -21 | 4 | -39 | -39 | 53 |
| -16 | -19 | 8 | -36 | -39 | 52 |
| -13 | -20 | 4 | -37 | -39 | 44 |
| -17 | -20 | 10 | -41 | -39 | 52 |
| -10 | -22 | 8 | -35 | -39 | 45 |
| -16 | -17 | 14 | -34 | -39 | 48 |
| -17 | -20 | 7 | -39 | -43 | 52 |
| -16 | -14 | 8 | -32 | -47 | 49 |
| -16 | -21 | 3 | -43 | -47 | 52 |
| -18 | -24 | 11 | -38 | -48 | 55 |
| -18 | -16 | 9 | -36 | -42 | 45 |
| -18 | -18 | 14 | -35 | -44 | 42 |
| -13 | -24 | 3 | -38 | -39 | 50 |
| -14 | -26 | 9 | -37 | -51 | 44 |
| -15 | -21 | 12 | -34 | -47 | 43 |
| -19 | -20 | 9 | -43 | -42 | 54 |
| -15 | -19 | 4 | -40 | -47 | 55 |
| -11 | -14 | 7 | -45 | -47 | 44 |
| -6 | -19 | 5 | -40 | -42 | 43 |

**Table S2: Peak coordinates of the right ROIs across subjects.**

|  | Thalamus |  |  | IPS |  |
| --- | --- | --- | --- | --- | --- |
| x | y | z | x | y | z |
| 14 | -24 | 14 | 42 | -35 | 51 |
| 13 | -11 | 12 | 39 | -39 | 56 |
| 19 | -22 | 11 | 40 | -45 | 48 |
| 15 | -11 | 7 | 36 | -40 | 54 |
| 13 | -16 | 8 | 47 | -41 | 51 |
| 11 | -17 | 2 | 48 | -44 | 52 |
| 10 | -21 | 9 | 48 | -43 | 51 |
| 9 | -13 | 5 | 44 | -47 | 44 |
| 10 | -17 | 2 | 45 | -39 | 48 |
| 18 | -20 | 10 | 39 | -44 | 53 |
| 11 | -19 | 7 | 36 | -37 | 47 |
| 15 | -20 | 8 | 45 | -39 | 43 |
| 19 | -22 | 8 | 43 | -35 | 52 |
| 7 | -19 | 4 | 39 | -42 | 42 |
| 18 | -16 | 7 | 34 | -41 | 47 |
| 11 | -19 | 9 | 46 | -46 | 49 |
| 17 | -17 | 10 | 45 | -39 | 47 |
| 17 | -14 | 8 | 47 | -38 | 46 |
| 15 | -20 | 7 | 46 | -45 | 46 |

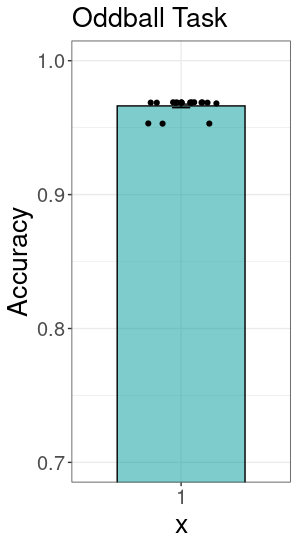

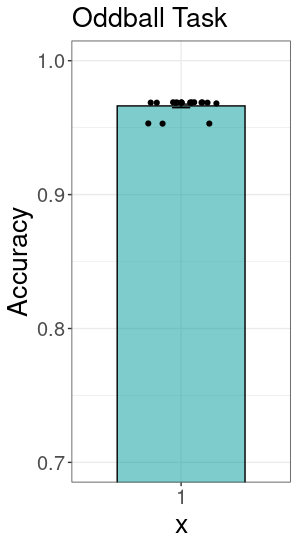

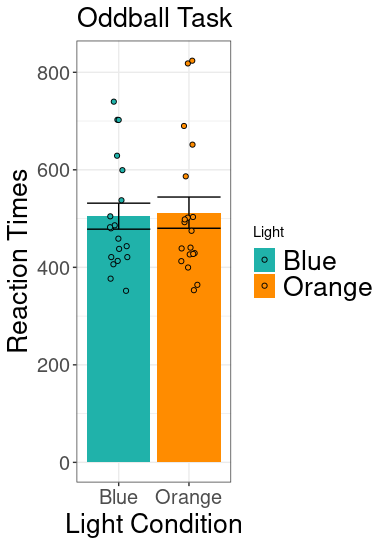

**A**

**B**

**Figure S1: Performance during the oddball task.** Participants were performing an acoustic version of the oddball task where they had to detect, by pressing a button with their right index, rare deviant tones among most frequent standard ones. While engaged in the task, they were exposed to short periods of blue-enriched or orange monochromatic light. A. Accuracy. As expected, participants’ accuracy (proportion of correct responses) was quite high (mean 0.96 ± 0.005) as shown here under. Participants’ individual performance is also shown. Y-axis limits were set from 0.7 to 1 to make individual data points more visible as they all lay at the top, outlining an almost ceiling performance. Accuracy is shown collapsed across light conditions. B. Reaction Times (RTs). RTs results are shown as a function of the light condition (Blue-enriched or Orange). Two dependent sample t-test: (*t*(19)= 2.10, *p*= 0.47, Cohen’s *d*= 0.05) did not show any significance of the light condition on the RTs. Individual data are shown. Error bars represent ±1 within-subjects Standard Error of the Mean (SEM) (Cousineau, 2005).
